# A Three-View System for CRISPR-Cas12a crRNA Activity Ranking: When to Ensemble, When Context Matters More

**DOI:** 10.64898/2026.11.07.765023

**Authors:** Tianze Deng, Zibo Lin, Hoyan Lam, Shirui Xu

## Abstract

CRISPR-Cas12a assays require effective crRNA selection, but experimentally screening all possible candidates is costly. We developed TriCasRank, a ranking model that combines sequence features with EasyDesign reference mapping and activity summaries from related training records. Historical summaries were calculated within each training fold to avoid information leakage. In five-fold target-grouped cross-validation using out-of-fold predictions, TriCasRank improved the Spearman correlation from 0.741 for the sequence-only ensemble to 0.821 (paired difference, 0.080; 95% target-group bootstrap interval, 0.072 to 0.090). However, its performance was comparable to that of a strong sequence-plus-mapping XGBoost model (SCC, 0.822; paired difference, -0.001; 95% interval, -0.007 to 0.005). Separate perturbation analyses supported the predictive value of mapping and historical features, although their effect sizes were not directly comparable because they were evaluated using different model configurations. Prediction errors remained larger at the extremes of the activity distribution. Performance also became less stable when template identities were separated between training and evaluation. TriCasRank may therefore be useful for ranking candidates represented within the EasyDesign-derived domain, but evaluation on an independent experimental cohort is still needed.

## 1. Introduction

### 1.1 Cas12a Diagnostics and crRNA Selection

CRISPR-Cas12a is a programmable RNA-guided DNA-targeting system. In molecular diagnostics, Cas12a can function as a molecular sensor: upon recognition of a complementary DNA target, the Cas12a protein activates a nonspecific single-stranded DNase activity, cleaving a fluorescent reporter and producing a measurable signal. This property makes Cas12a a promising platform for nucleic acid diagnostics [1].

Selecting effective crRNAs remains a critical constraint in assay design. For a given target sequence, hundreds of candidate crRNAs can be designed. Thus, testing every candidates is costly and time-consuming, particularly when rapid assay development is required. Machine-learning models can help prioritize candidates for experimental validation [2,3].

Mismatch count is not the only thing that determines Cas12a activity.

Other factors such as Mismatch position and identity, nucleotide composition, and local sequence context can also influence guide activity. These properties form the basis of many sequence-based prediction models [3].

### 1.2 Sequence Features and Contextual Information

Most sequence-based predictors represent each crRNA–target pair using features such as mismatches, nucleotide composition, positional identity, or local sequence patterns. These representations describe the current pair but may not capture relationships recorded in the experimental reference library. In the EasyDesign dataset, a target may also be associated with a reference template, template group, mapping orientation, and mapping status. Such variables provide information about the target’s position within the source library. Sequence-based Cas12a activity models have used related guide, target, and positional representations [2,3].

Repeated guide and template identities provide another potential source of predictive information. Activity measurements from related training records can be summarized as guide- or template-level historical features. However, these summaries must be calculated without using information from held-out records; otherwise, performance estimates may be affected by data leakage. Their value should also be distinguished from the simpler effect of increasing the number of input variables [4].

Model ensembling offers a separate route to improved prediction. Combining models may reduce error when their predictions are sufficiently accurate and their errors are not identical. However, the benefit of adding another model should be evaluated against the benefit of adding a new source of information. In this study, we therefore examined contextual features and model fusion as related but distinct components of crRNA activity prediction [5].

### 1.3 Study Objectives

We developed TriCasRank, a three-module system for ranking Cas12a crRNA–target activity. Module D uses sequence-derived features, whereas Modules B and C incorporate reference-library mapping and training-derived guide and template information. The fixed-weight TriCasRank configuration was developed using target-grouped out-of-fold predictions. Nested cross-validation was subsequently used to evaluate model selection without reusing the outer-fold outcomes.

The study examined whether mapping and historical information improved activity ranking beyond sequence features alone. We also evaluated the contribution of each module, the effect of different identity-grouping schemes, and performance relative to a locally retrained EasyDesign model on a matched cohort.

Because guide and template identities were highly represented across the development and historical evaluation data, the main results describe performance within the EasyDesign-derived domain. Evaluation under stricter identity separation was used to assess how performance changed when contextual support became less available.

## 2. Materials and Methods

### 2.1 Data and Scope

We analyzed 10,634 fluorescence-assay records reported with EasyDesign [2]. We retained the historical partition established during earlier model development. Of the 10,634 records, 8,417 were used for model development and 2,217 were used as a fixed-validation set. The earlier partition was generated deterministically from the normalized target sequence. The first eight hexadecimal characters of its SHA-1 digest were converted to an integer, and records with a value modulo 100 below 80 were assigned to development; the remaining records were assigned to historical validation. This rule reproduced all 10,634 assignments. For every record, the available inputs were a 25-position crRNA, its aligned 25-position target, and a processed activity label derived from the source study’s 30-min fluorescence readout. We used the published Table S3 30-min activity values without further label processing. The source study describes plate-level normalization using negative and high-activity positive controls, but the sample-level raw readings and plate identifiers were unavailable, so the complete raw-to-label transformation could not be independently reconstructed. The separate 20-min normalized field and Table S4 augmentation were not used. MAE and RMSE are on the published activity-label scale [2].

The sequences were already aligned to 25 positions before they entered the pipeline. We treated the 30-min endpoint as the response and did not model the reaction trajectory. One feature of the data is especially important for interpretation: 95.4% of records in the historical validation set (2,116/2,217) contained a guide sequence that also occurred in the development set. At the unique-guide level, the corresponding overlap was 992/1,008 (98.4%). The primary analysis consequently measures performance largely within the EasyDesign-compatible guide distribution. Section 2.8 reports the additional identity-isolated analyses.

### 2.2 Feature Engineering

We described each sample from three sources of information: the aligned sequence pair, its source context, and activity patterns available from the training data. The original sequence set contained 188 candidate variables. Five were constant in the training partition, so 183 sequence/alignment features remained for modeling. The nucleotide pair at each aligned position was one-hot encoded to distinguish substitution types at the same position.

To derive mapping features, target sequences were matched against the frozen EasyDesign Table S2 reference [2] in both forward and reverse-complement orientations. Exact sequence-window matches were prioritized, and IUPAC-compatible matching was used only when no exact match was available. Historical activity summaries for recurring guides and template keys were recomputed separately within each outer training fold and then applied to the corresponding held-out fold. Thus, activity labels from the held-out fold were not used to construct these summaries.

**Table 1.** Feature blocks organized by information view.

| **View** | **Feature block** | **Count** | **Description** |
|---|---|---|---|
| Sequence pair | Active sequence features | 183 | Mismatch, gap, GC, composition and local sequence-context features |
| Sequence pair | Position-specific pair identities | 453 | Encoded nucleotide pairing at aligned positions |
| Historical prior | Guide statistics | 11 | Smoothed training-set statistics for recurring guides |
| Historical prior | Template-key statistics | 12 | Smoothed training-set statistics for mapped template keys |
| Source context | Mapping numeric counts | 4 | Candidate template hits, groups and orientations |
| Source context | Mapping category indicators | 528 | Template identity, group identity and mapping status |

*Note. The complete mapping-aware representation contained 1,191 columns, including one-hot-encoded categorical variables; the column count should therefore not be interpreted as the number of independent biological factors. A template key denotes the retained EasyDesign reference-template identifier or, where applicable, a deterministic combination of compatible candidate identifiers.*

### 2.3 Validation Protocols

We used four complementary evaluation procedures: target-grouped out-of-fold evaluation, historical fixed validation, identity-grouped evaluation, and nested cross-validation. Model development relied on five-fold target-grouped OOF validation of the 8,417 training records, with identical target sequences kept in the same fold. The same predefined fold assignments were used for development-stage architecture comparisons and fixed-weight searches. In the nested analysis, model and weight selection were confined to the inner folds, and performance was evaluated on the corresponding outer fold. The 2,217-record historical fixed-validation set was then used to compare trained models. Since this set had already been inspected during earlier development, we did not regard it as a fully untouched external test set. Uncertainty was estimated by target-cluster bootstrap so that records from the same target stayed together during resampling [6–8].

**Table 2.** Validation procedures and their purposes.

| **Protocol** | **Records** | **Purpose** |
|---|---|---|
| Historical fixed validation | 2,217 | Comparison of trained models on a fixed historical partition |
| Target-grouped OOF | 8,417; 5 folds | Model development, architecture comparison and weight selection |
| Target-cluster bootstrap | 2,000 iterations | Uncertainty estimation with target-level resampling |

*Note. The three procedures served different purposes, and their estimates were not treated as directly interchangeable.*

Spearman’s rank correlation coefficient (SCC) was the primary evaluation metric. Pearson’s correlation coefficient (PCC), root mean squared error (RMSE), mean absolute error (MAE), and R² were reported as secondary metrics.

### 2.4 TriCasRank Architecture: Three-Module Ensemble

TriCasRank combines predictions from three modules representing sequence, mapping context, and historical context. The final prediction was calculated as:

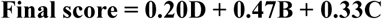

Module D uses sequence information only. It takes the 183 active sequence features and combines XGBoost [9], LightGBM [10], and MLP predictions with OOF-selected weights of 0.35, 0.59, and 0.06. Modules B and C share the full 1,191-column mapping-aware input but use different historical anchors. Module B uses the smoothed guide-level activity mean as its anchor, whereas Module C uses an equally weighted combination of the smoothed guide-level and template-key means. In both modules, XGBoost predicts a residual correction to the corresponding sample-specific anchor. When label-free reference mapping is unavailable, prediction uses the sequence-only Module D. An unseen historical key uses the fitted history fallback within contextual feature calculation.

### 2.5 Weight Search and Model Selection

Fixed D/B/C weights were determined via grid search on target-grouped out-of-fold predictions using 0.005 increments, subject to a sum-of-one constraint. Combinations yielding an SCC within 0.0001 of the maximum were retained to assess stability. Since the same out-of-fold predictions were used for both model development and weight optimization, this score serves as a development estimate. To validate robustness, a separate nested cross-validation was performed, repeating the model and weight selection within the inner folds and evaluating the final configuration on the outer folds. For each of five outer folds, the other four folds provided four training-only inner validation folds. D-component weights were selected first, followed by D+B and D/B/C weights, then the model family. The step-0.05 grids included the original fixed weights. The selected rule was evaluated only on its outer query fold. SELECTED_POLICY was the primary nested comparison; nested TriCasRank architecture effects were reported separately.

### 2.6 Heterogeneous Ensembling Experiments

Development-stage ensemble observations included the sequence-only XGBoost–LightGBM–MLP ensemble and the transition from the legacy A–B–C system to the complete D/B/C system. The sequence-only result is reported under fixed target-grouped evaluation, whereas the legacy-system comparison uses archived cross-fitted meta-OOF predictions. These estimates are not directly interchangeable.

### 2.7 Controlled Information-Block and Perturbation Analyses

We used ablation and permutation experiments to determine whether context added information rather than merely increasing the number of input columns. The learner was kept as centered-target XGBoost while its inputs were changed among four conditions: sequence only, sequence with history, sequence with mapping, and all three blocks. The two perturbation controls were applied to different predictors. Historical labels were shuffled within each outer training fold for the full TriCasRank model, whereas the mapping block was permuted for the sequence-plus-mapping XGBoost model. Each perturbed model was compared with its own intact control; therefore, the effect sizes from the two experiments were not directly comparable. Sequences, fold membership, held-out labels, and model capacity otherwise remained fixed. Each perturbation was repeated with three predefined seeds, with uncertainty estimated from 2,000 paired target-group bootstrap samples.

### 2.8 Identity-Isolated Evaluation

To examine how performance changed under stricter separation, we repeated OOF evaluation using different grouping identities [6]. The four settings were target group, crRNA sequence, atomic template, and a joint target-template-guide identity, all evaluated on the same 8,417 records. TriCasRank was compared with Module D, the history-prior baseline, the sequence-plus-mapping XGBoost model, and the fixed D+B ensemble using SCC and error metrics, with paired confidence intervals based on resampling the relevant groups. We also calculated within-guide SCC for guides represented by at least five records, which measures how well targets can be ranked for the same crRNA. All historical features were recomputed within each training fold under every grouping scheme.

### 2.9 Matched Comparison with EasyDesign

For a direct comparison with EasyDesign [2], we used only records that could be represented consistently by both methods. Of the 8,417 development records, 5,995 had complete and unambiguous inputs for both methods and formed the matched cohort, representing 5,923 target groups. The remaining records were excluded because of ambiguous mappings (n = 2,279), incomplete contextual inputs (n = 142), or absence of a mapping (n = 1). We locally retrained the released EasyDesign regression model on the shared cohort using the same target-grouped folds, giving the EasyDesign-adapted reference used in our comparison. Target-grouped cross-fitted predictions for EasyDesign-adapted, TriCasRank, and Module D were generated on the same records. We compared SCC, RMSE, paired lower-error frequency, and MAE across activity quintiles, with 2,000 target-group bootstrap resamples for uncertainty. The published EasyDesign score was cited only as background and was not used as a same-record benchmark.

*Note. EasyDesign-adapted refers to* local *retraining of the released EasyDesign regressor on the shared cohort, not to the published EasyDesign test score*.

## 3. Results

### 3.1 Overall Benchmark Performance

All models were evaluated on the same 8,417 records from 6,825 target groups using five-fold target-grouped cross-validation. Records sharing the same target identity were assigned to a single fold, and the held-out predictions from all folds were pooled for evaluation (Fig. 1a).

**Fig. 1.**
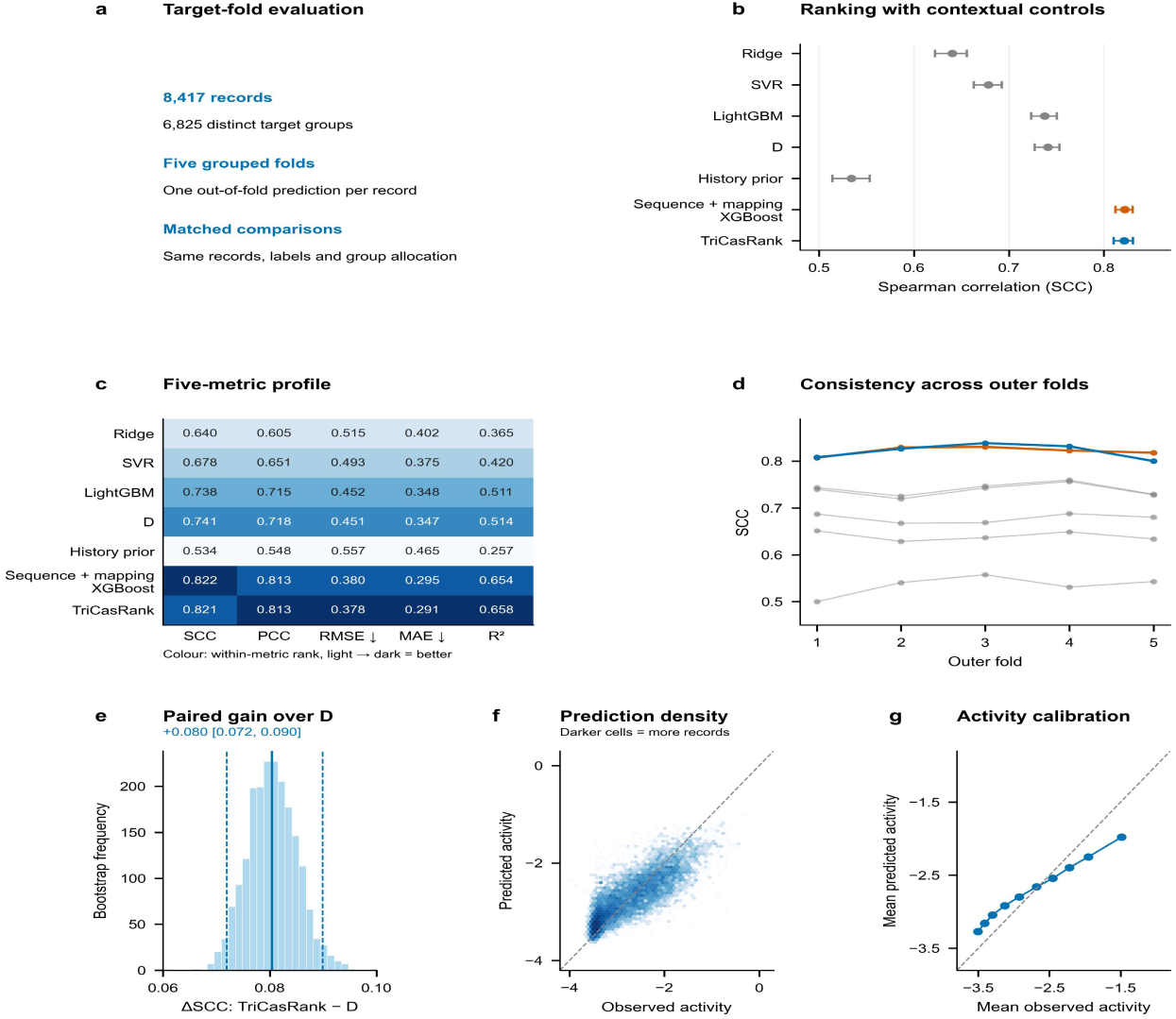
Context-aware prediction under target-grouped evaluation. a, Shared 8,417-record, five-fold protocol with 6,825 target groups. b, SCC for seven models; points and bars show pooled OOF estimates and 95% target-group bootstrap intervals. c, Five-metric profiles; numbers are raw values and colour is the direction-corrected within-metric rank. d, Outer-fold SCC, with the same colour key as b: TriCasRank blue, sequence-plus-mapping XGBoost orange, other controls grey. e, Paired bootstrap distribution for TriCasRank − D. f, Prediction density using all records; darker cells contain more records. g, Calibration in ten observed-activity bins. Dashed lines in f,g denote identity. Bootstrap analyses use 2,000 target-group samples, seed 20261012.

We compared Ridge, SVR, LightGBM, the sequence-only Module D, a history-prior reference, TriCasRank, and a strong sequence-plus-mapping XGBoost model. TriCasRank increased SCC from 0.741 for Module D to 0.821, a gain of 0.080 (95% target-group bootstrap interval, 0.072–0.090). However, the sequence-plus-mapping XGBoost model achieved a similar SCC of 0.822, and the difference relative to TriCasRank was −0.001 (95% interval, −0.007 to 0.005). TriCasRank showed slightly lower RMSE (0.378 versus 0.380) and MAE (0.291 versus 0.295) (Table 3). In nested cross-validation, the sequence-plus-mapping XGBoost model was selected in 37 of 40 outer-fold evaluations, whereas TriCasRank was selected in 3, indicating that the additional historical fusion did not consistently improve ranking beyond the strong mapping-aware model. The development-stage observations in Table 4 document the sequence-only ensemble and the transition from the legacy A–B–C system to the complete D/B/C system; they use separate evaluation protocols.

**Table 3.** Performance of TriCasRank and reference models under target-grouped cross-validation.

| **Model** | **Role and input** | **Records** | **Groups** | **SCC (95% CI)** | **PCC** | **RMSE** | **MAE** | **R²** |
|---|---|---|---|---|---|---|---|---|
| Ridge | Direct sequence baseline; 183 active sequence/alignment features | 8417 | 6825 | 0.640 (0.622–0.656) | 0.605 | 0.515 | 0.402 | 0.365 |
| SVR | Direct sequence baseline; 183 active sequence/alignment features | 8417 | 6825 | 0.678 (0.663–0.692) | 0.651 | 0.493 | 0.375 | 0.420 |
| LightGBM | Direct sequence baseline; 183 active sequence/alignment features | 8417 | 6825 | 0.738 (0.723–0.750) | 0.715 | 0.452 | 0.348 | 0.511 |
| D | Sequence-only ensemble; 0.35 XGBoost + 0.59 LightGBM + 0.06 MLP | 8417 | 6825 | 0.741 (0.727–0.753) | 0.718 | 0.451 | 0.347 | 0.514 |
| History prior | Training-fold historical-prior reference | 8417 | 6825 | 0.534 (0.514–0.553) | 0.548 | 0.557 | 0.465 | 0.257 |
| Sequence + mapping XGBoost | Matched mapping-aware single learner | 8417 | 6825 | 0.822 (0.812–0.830) | 0.813 | 0.380 | 0.295 | 0.654 |
| TriCasRank | Three-view system: sequence-only D, mapping-aware B, and history-anchored C; 0.20/0.47/0.33 with D fallback | 8417 | 6825 | 0.821 (0.810–0.831) | 0.813 | 0.378 | 0.291 | 0.658 |

*Note. All metrics were calculated from pooled out-of-fold predictions for the same 8,417 records and 6,825 target groups. Values in parentheses are 95% intervals obtained from 2,000 target-group bootstrap resamples. Higher SCC, PCC, and R² indicate better performance, whereas lower RMSE and MAE indicate better performance.*

**Table 4.** Development-stage ensemble comparisons and evaluation protocols.

| **Development observation** | **Evaluation** | **Verified result** |
|---|---|---|
| Sequence-only Module D | Current fixed target-grouped OOF | SCC 0.741 |
| Complete DBC replacing legacy ABC | Earlier cross-fitted meta OOF | ΔSCC +0.00236; 95% CI [+0.00106, +0.00365] |

*Note. The two rows use different evaluation protocols. The archived legacy-system comparison is a development observation, not a current fixed target-grouped result.*

Predictions retained a clear monotonic relationship with observed activity but were compressed toward the center of the activity range (Fig. 1f,g). Thus, TriCasRank improved relative ranking more strongly than absolute calibration at the activity extremes.

### 3.2 Contribution of Contextual Information

TriCasRank combines the sequence-only Module D with two mapping-aware residual modules, B and C, using fixed weights of 0.20D + 0.47B + 0.33C. Modules B and C share the same mapping-aware inputs but use different historical anchors: Module B uses the guide-level activity mean, whereas Module C uses an equally weighted combination of the guide-level and template-key means. When the required historical anchor is unavailable, the corresponding contextual prediction falls back to Module D (Fig. 2a).

**Fig. 2.**
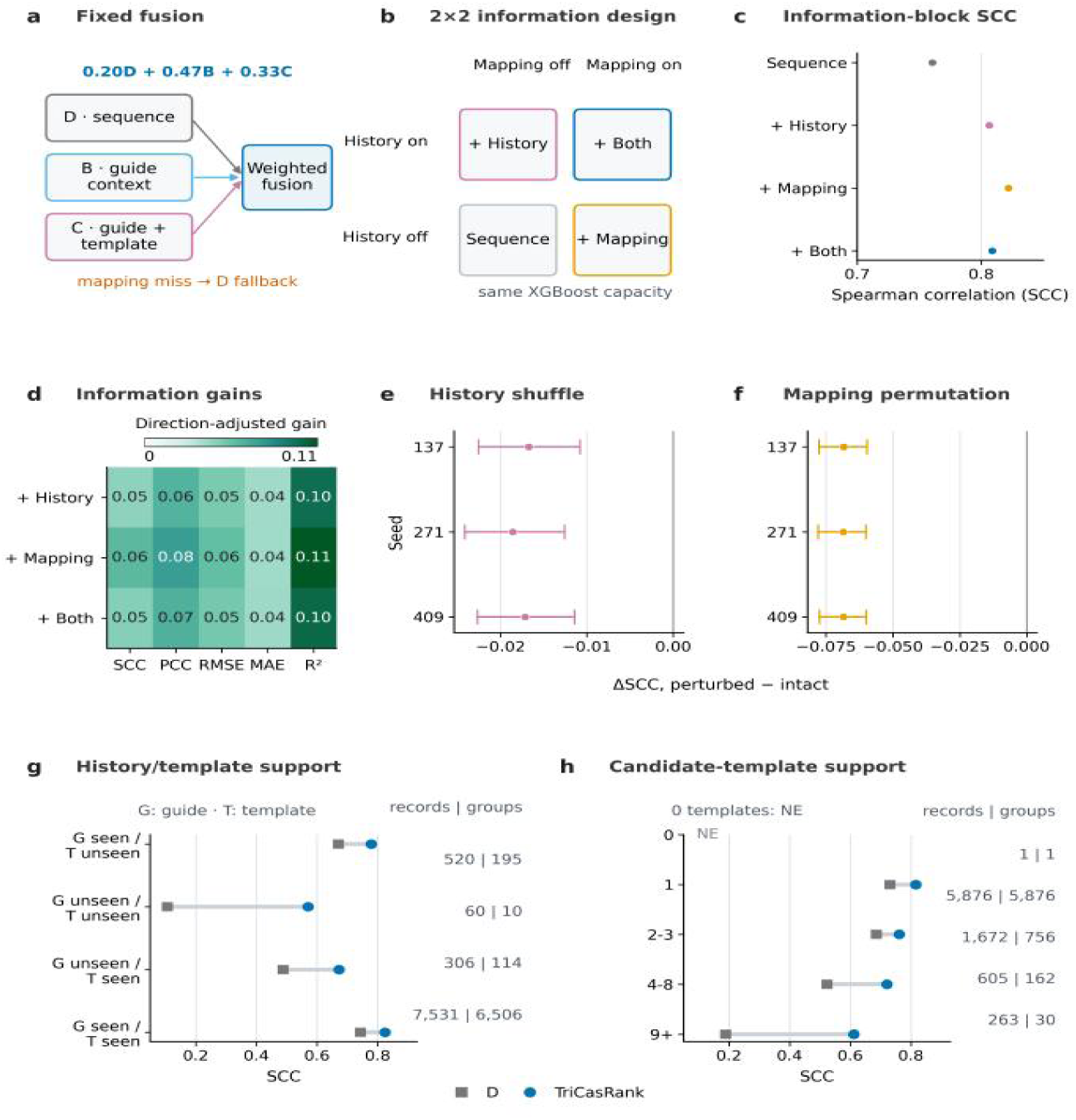
Contribution of mapping and historical-context information to TriCasRank performance. **a,** TriCasRank combines Module D with mapping-aware residual Modules B and C using fixed weights of 0.20D + 0.47B + 0.33C. Module B uses a guide-level historical anchor, whereas Module C uses combined guide- and template-level anchors; unavailable anchors trigger fallback to Module D. Records without the required contextual support use the sequence-only Module D prediction. **b,** Matched-capacity 2×2 design that varies history and mapping while holding the centred-target XGBoost learner fixed. **c,** SCC for sequence only, sequence + history, sequence + mapping and sequence + history + mapping on 8,417 records from 6,825 target groups. **d,** Five-metric changes relative to sequence only; Values were direction-adjusted, with positive values indicating improvement. e, Change in SCC after shuffling the training labels used to construct historical summaries in the full TriCasRank model. f, Change in SCC after permutation of the mapping block in the sequence-plus-mapping XGBoost model; the magnitudes in e and f are not directly comparable. **g,** Performance of TriCasRank and Module D according to the availability of guide- and template-key historical support. **h,** Performance according to candidate-template support. SCC was not calculated for the stratum containing only one record.

A matched-capacity 2×2 design held the centred-target XGBoost learner fixed while switching history and mapping blocks on or off, converting the architectural question into a controlled test of information content (Fig. 2b). SCC increased from 0.760 with sequence alone to 0.806 with history and to 0.822 with mapping, whereas the combined history-plus-mapping arm reached 0.809, placing explicit guide–template correspondence at the top of the controlled comparison (Fig. 2c). Direction-adjusted changes in all five metrics confirmed that mapping improved both ordering and numerical accuracy relative to sequence, while the combined arm showed that the two contextual blocks contributed overlapping rather than simply additive information (Fig. 2d).

Two separate perturbation analyses evaluated the association between contextual information and predictive performance. In the full TriCasRank model, shuffling the training labels used to construct historical summaries reduced SCC by 0.017–0.019 across three predefined seeds. In the sequence-plus-mapping XGBoost model, permuting the mapping block reduced SCC by approximately 0.068. Because the two perturbations were applied to different predictors, their effect sizes should not be used to rank the relative importance of mapping and historical information.

Performance was then examined according to the availability of guide- and template-level historical support. Most records—7,531 records from 6,506 target groups—had both types of support (Fig. 2g). As contextual support decreased, performance estimates became less stable because the corresponding strata contained fewer records. These support categories should not be interpreted as direct counts of Module D fallback events., and estimates for the smallest strata were correspondingly less stable (Fig. 2g,h).

### 3.3 Local specialization converts correlated views into a stronger fusion

Modules B and C produced highly correlated predictions (r=0.990) and residuals (r=0.982), whereas Module D showed lower residual correlations with the two contextual modules (r=0.765–0.784; Fig. 3a–c). This indicates that B and C shared most global error structure while D supplied a different error profile. Despite the high correlation between B and C, the module with the smallest absolute error varied across records. Module D, B, and C uniquely produced the lowest absolute error for 35.4%, 34.2%, and 30.3% of records (Fig. 3d).

**Fig. 3.**
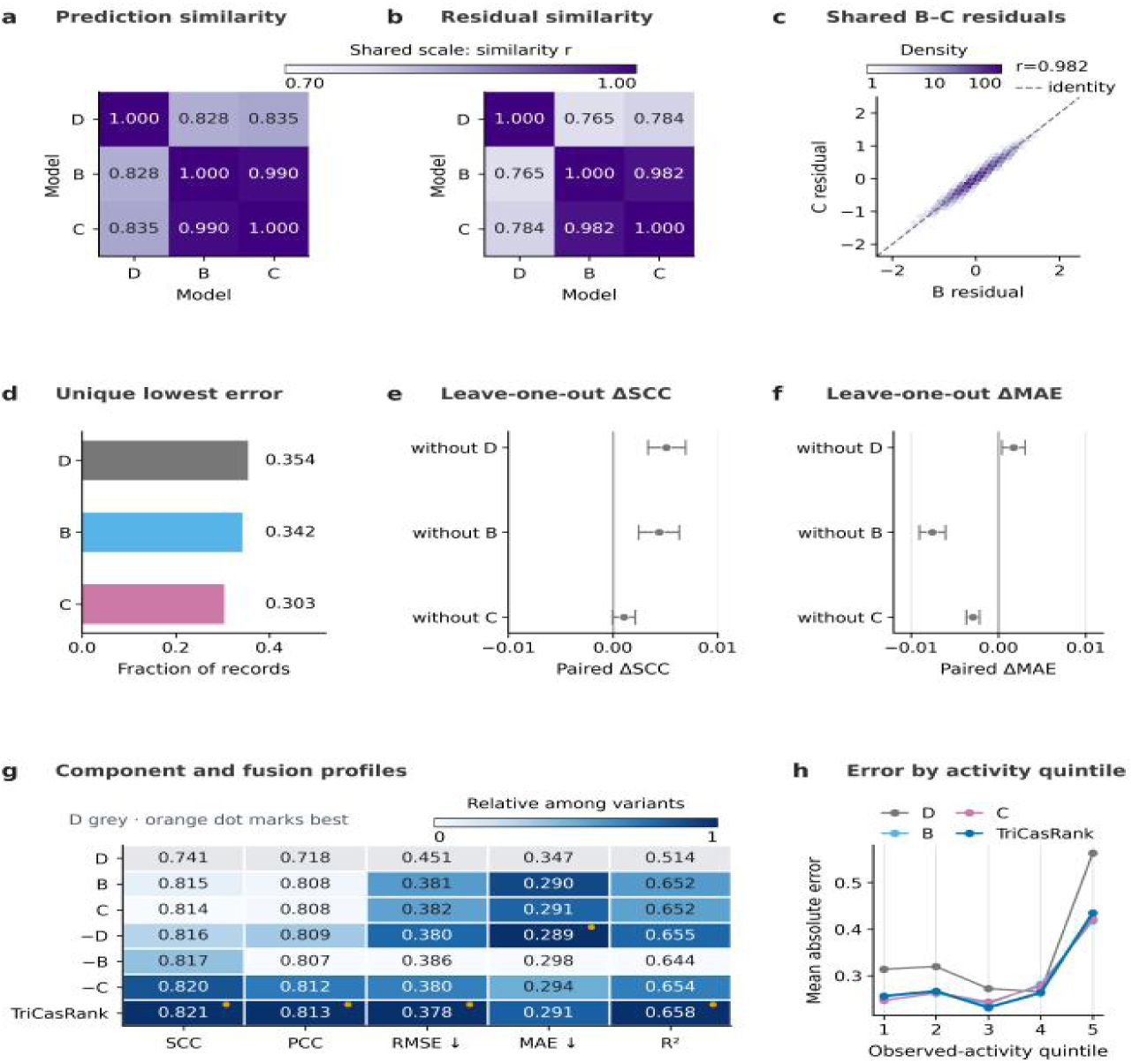
Prediction dependence and component-removal analysis for Modules D, B, and C. **a,** Pairwise prediction correlations for D, B and C. **b,** Pairwise residual correlations on the same 0.70–1.00 scale; both matrices use 8,417 records. **c,** Hexbin density of B and C residuals, with identity shown by the dashed line. **d,** Fraction of records uniquely predicted with the lowest absolute error by D, B or C. **e,** Paired SCC change after removing one component from the fixed-weight fusion and renormalizing the remaining weights. **f,** Corresponding MAE change. Intervals in e and f are 95% intervals from 2,000 paired bootstrap resamples of 6,825 target groups; the same out-of-fold component predictions are reused. **g,** Five-metric profiles for the components, component-removal variants and TriCasRank. Cell text gives raw values, colour gives direction-corrected within-metric performance and orange dots mark the best displayed value. **h,** Mean absolute error across five observed-activity quintiles, each containing 1,683 or 1,684 records. The panels summarize prediction dependence, component-removal effects, and activity-dependent error patterns across the three modules.

We assessed each module’s contribution by removing one component from the fixed-weight fusion and renormalizing the remaining weights. Removing Module D reduced SCC by approximately 0.005 but slightly improved MAE. Removing Module B or C had smaller effects on SCC but increased MAE (Fig. 3e,f; Table 5). Thus, Module D contributed more strongly to rank performance in this comparison, whereas the contextual modules contributed more to absolute-error reduction.

**Table 5.** Performance of individual modules, feature-block configurations, and component-removal variants.

| Block | System | Information | Role or weight | SCC | PCC | RMSE | MAE | R2 | Paired contrast |
|---|---|---|---|---|---|---|---|---|---|
| Components / endpoint | D | 183 active sequence/alignment features | Sequence-only reference; fallback view | 0.741 | 0.718 | 0.451 | 0.347 | 0.514 | - |
| Components / endpoint | B | Mapping-aware contextual features | Context view; fixed final weight 0.47 | 0.815 | 0.808 | 0.381 | 0.290 | 0.652 | - |
| Components / endpoint | C | Guide/template historical anchors with mapping-aware residual features | Context view; fixed final weight 0.33 | 0.814 | 0.808 | 0.382 | 0.291 | 0.652 | - |
| Components / endpoint | TriCasRank | D + B + C relational views | 0.20D + 0.47B + 0.33C; D fallback | 0.821 | 0.813 | 0.378 | 0.291 | 0.658 | - |
| Matched-capacity information blocks | Sequence only | Sequence block | Centered-target XGBoost; capacity held fixed | 0.760 | 0.737 | 0.439 | 0.336 | 0.540 | Reference arm |
| Matched-capacity information blocks | Sequence + history | Sequence + history block | Centered-target XGBoost; capacity held fixed | 0.806 | 0.801 | 0.387 | 0.295 | 0.641 | Reference arm |
| Matched-capacity information blocks | Sequence + mapping | Sequence + mapping block | Centered-target XGBoost; capacity held fixed | 0.822 | 0.813 | 0.380 | 0.295 | 0.654 | Reference arm |
| Matched-capacity information blocks | Sequence + history + mapping | Sequence + history + mapping blocks | Centered-target XGBoost; capacity held fixed | 0.809 | 0.803 | 0.386 | 0.294 | 0.644 | Reference arm |
| Fixed-weight fusion / removal | D–B–C unrouted | D + B + C component predictions | Frozen 0.20/0.47/0.33 weights without fallback routing | 0.821 | 0.813 | 0.378 | 0.291 | 0.658 | Reference for removal contrasts |
| Fixed-weight fusion / removal | Remove D | B + C component predictions | Same-routing fusion; remaining weights renormalized | 0.816 | 0.809 | 0.380 | 0.289 | 0.655 | ΔSCC 0.005099 [0.003328, 0.006905]; ΔMAE 0.001718 [0.000373, 0.003050] |
| Fixed-weight fusion / removal | Remove B | D + C component predictions | Same-routing fusion; remaining weights renormalized | 0.817 | 0.807 | 0.386 | 0.298 | 0.644 | ΔSCC 0.004417 [0.002435, 0.006302]; ΔMAE −0.007577 [−0.009068, −0.006079] |
| Fixed-weight fusion / removal | Remove C | D + B component predictions | Same-routing fusion; remaining weights renormalized | 0.820 | 0.812 | 0.380 | 0.294 | 0.654 | ΔSCC 0.001049 [−0.000040, 0.002100]; ΔMAE −0.002935 [−0.003673, −0.002184] |

*Note. All values were calculated from pooled target-grouped out-of-fold predictions. For component-removal analyses, the remaining fixed weights were renormalized to sum to one. ΔSCC and ΔMAE were calculated as the performance of the full D–B–C fusion minus that of the corresponding removal variant. Values in brackets are 95% paired target-group bootstrap intervals.*

The final TriCasRank model achieved the best SCC, PCC, RMSE, and R² among the displayed configurations, whereas the model without Module D achieved a lower MAE (Fig. 3g; Table 5). The quintile analysis helps explain this difference. Fusion reduced much of Module D’s error in the high-activity range, although predictions remained compressed near both ends of the scale (Fig. 3h). Thus, the three-view design succeeds through local specialization rather than wholesale disagreement: D preserves rank-sensitive structure, B and C refine magnitude, and their fixed fusion converts those roles into the strongest overall ranking profile (Fig. 3d–h).

### 3.4 TriCasRank retains ranking under identity shift

We repeated the OOF evaluation using four grouping schemes based on target identity, crRNA sequence, atomic-template membership, or joint target–template–guide membership (Fig. 4a,b). The same 8,417 records were retained in each analysis, while the unit assigned to a fold differed among the protocols.

**Fig. 4.**
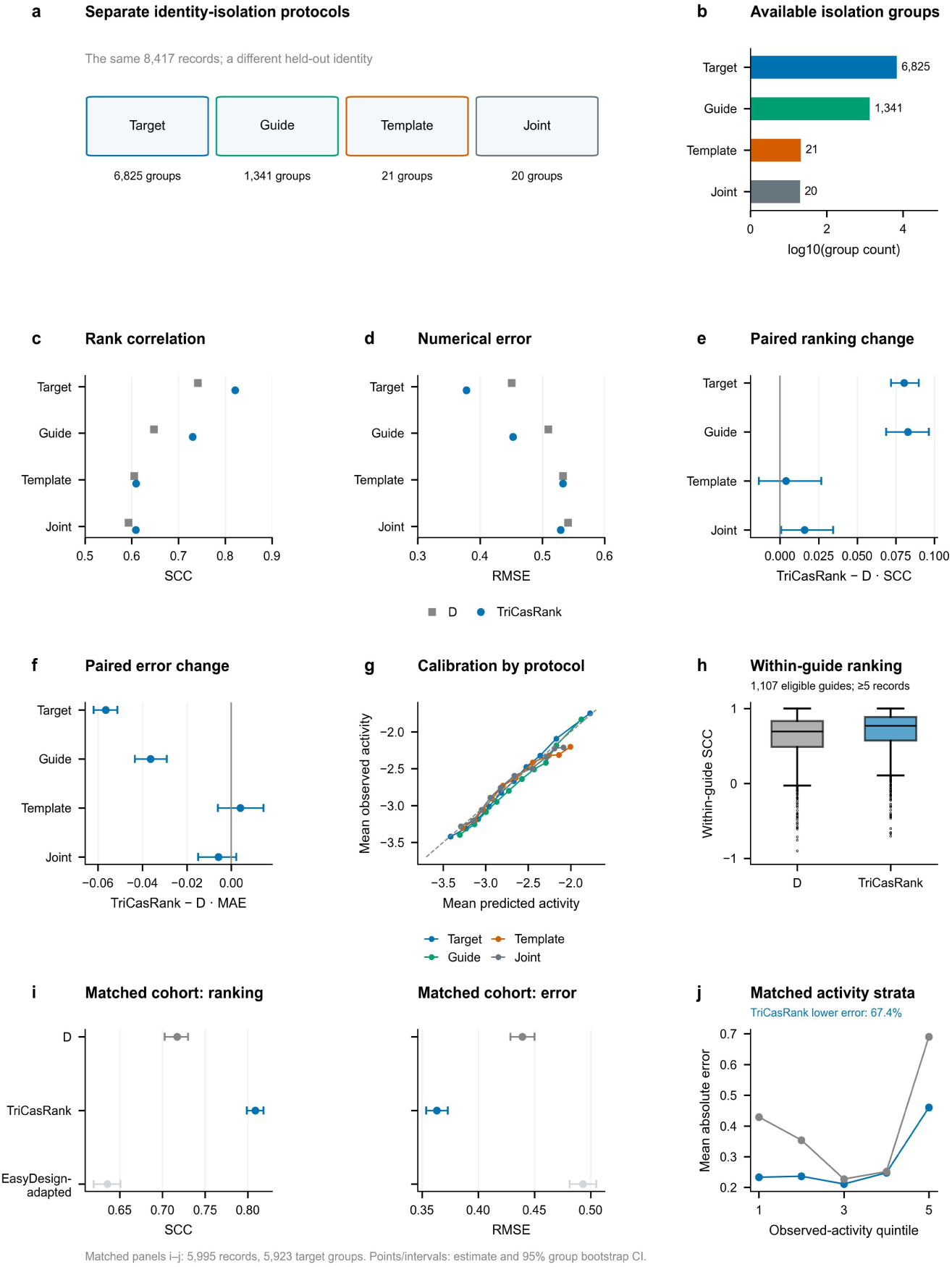
Identity-dependent transfer and the matched EasyDesign comparison. a,b, Four separate identity-isolation protocols and their group counts, using the same 8,417 records. c,d, Pooled OOF SCC and RMSE for D and TriCasRank. e,f, Paired differences, TriCasRank − D, with 95% intervals from 2,000 samples of the corresponding isolation groups. g, Calibration in ten prediction-ordered bins. h, Guide-isolated within-guide SCC for 1,107 guides with at least five records; boxes show median and interquartile range, whiskers use 1.5× IQR, and points show outliers. i, SCC and RMSE with 95% target-group bootstrap intervals on the matched 5,995 records. j, MAE by observed-activity quintile; TriCasRank has lower absolute error on 67.4% of paired records. EasyDesign-adapted is a local retraining of the released regressor under the shared protocol.

Relative to Module D, TriCasRank increased SCC by 0.080 under target grouping and by 0.083 under guide grouping, while reducing MAE by 0.056 and 0.036, respectively. Under guide isolation, however, the sequence-plus-mapping XGBoost model achieved a higher SCC than TriCasRank (0.773 versus 0.730). Within 1,107 eligible guides, the median within-guide SCC increased from 0.696 for Module D to 0.771 for TriCasRank, indicating improved ranking of targets for the same crRNA.

Under atomic-template grouping, TriCasRank produced only a small SCC increase over Module D (0.609 versus 0.605) and a slightly higher MAE (0.419 versus 0.415). Under joint grouping, TriCasRank achieved an SCC of 0.609 compared with 0.593 for Module D, while MAE decreased slightly from 0.419 to 0.413. The sequence-plus-mapping XGBoost model performed better than TriCasRank under both stricter protocols, reaching SCC values of 0.659 and 0.651 for atomic-template and joint grouping, respectively. Thus, the fixed TriCasRank fusion did not provide the strongest transfer performance under stricter identity separation.

### 3.5 Matched Comparison with EasyDesign

We next compared TriCasRank with a locally retrained EasyDesign regressor using the same 5,995 records from 5,923 target groups. All models were trained and evaluated using the same target-grouped folds, avoiding direct comparison with the previously published EasyDesign test score (Fig. 4i).

TriCasRank achieved an SCC of 0.809 and an RMSE of 0.363, compared with 0.635 and 0.493, respectively, for the EasyDesign-adapted model. TriCasRank also produced a smaller absolute error for 67.4% of paired records and had lower MAE in each observed-activity quintile (Fig. 4i,j). These findings apply to the shared cohort and locally retrained EasyDesign implementation rather than to the original published benchmark.

## 4. Discussion

### 4.1 Principal Findings

TriCasRank improved ranking relative to the sequence-only Module D. However, its SCC was comparable to that of the sequence-plus-mapping XGBoost model, indicating that mapping context accounted for most of the observed ranking improvement, whereas the additional historical fusion provided limited incremental benefit under the primary protocol.In the target-grouped analysis, SCC increased from 0.741 for Module D to 0.821 for TriCasRank. PCC and R² also increased, RMSE and MAE decreased, and every outer fold showed a positive SCC difference (Fig. 1b–e; Table 3). These results apply to the LbCas12a assays and data structure evaluated here.

The controlled experiments provided separate evidence that both mapping correspondence and historical summaries contained predictive information. Permuting the mapping block reduced SCC by approximately 0.068 in the sequence-plus-mapping XGBoost model, whereas shuffling the labels used to construct historical summaries reduced SCC by 0.017–0.019 in the full TriCasRank model. Because these perturbations were applied to different predictors, their magnitudes cannot be compared directly or used to determine which contextual source was more influential.

The calibration plots also reveal a limitation: predicted values were drawn toward the center of the activity range. TriCasRank should consequently be viewed first as a ranker within the studied domain. Improving numerical calibration, particularly at the two extremes, remains a separate task (Fig. 1f,g; Fig. 3h).

### 4.2 Contribution of Contextual Information

Context helped, but the two context blocks did not produce independent gains. Sequence with mapping reached an SCC of 0.822, compared with 0.809 when both mapping and history were included (Fig. 2c,d; Table 5). Mapping and historical summaries may therefore encode some of the same relationships. This interpretation is also consistent with the perturbation results: breaking the mapping correspondence produced the larger loss, while shuffled historical labels produced a smaller but reproducible decline (Fig. 2e,f; Table S3).

However, these contextual variables should not be interpreted as direct measurements of a biological mechanism. Mapping identifiers may capture several correlated factors, including target family, assay composition, sequence similarity, or other dataset-specific properties. Similarly, the effect of historical features depends on the recurrence of guide and template identities between training and evaluation data. The present analysis shows that these variables improve prediction within the studied dataset, but they do not establish that guide–template correspondence is itself a causal determinant of Cas12a activity.

The support-stratified analysis was descriptive. Most records had both guide- and template-level support, whereas strata with limited support contained fewer observations and produced less stable estimates. When a required contextual key was unavailable, the model specification used Module D as a fallback. However, the support categories should not be interpreted as direct counts of fallback events or as evidence that contextual support caused the observed performance differences.

### 4.3 Module Complementarity and Calibration

The modules differed at the record level even when their overall predictions were similar. Modules B and C were strongly correlated in both predictions and residuals, whereas Module D had lower residual correlations with them (Fig. 3). Even so, no component dominated the record-wise error comparison: D, B, and C each supplied the unique lowest absolute error for roughly one third of the records.

Component-removal analyses suggested different contributions to the final prediction. Removing Module D reduced SCC but slightly improved MAE, whereas removing Module B or C had smaller effects on SCC but increased MAE. Within this particular fixed-weight configuration, Module D therefore contributed more to rank performance, while the contextual modules contributed more to absolute-error reduction. This interpretation is specific to the selected weights and evaluated data and should not be treated as a general functional division among model classes.

The full fusion achieved the best SCC, PCC, RMSE, and R² among the displayed configurations, while the model without Module D achieved a slightly lower MAE. That balance is acceptable for a system designed to rank candidates. Applications that depend on an activity threshold would need a separate calibration step, with particular attention to the ends of the activity range.

### 4.4 Performance Under Alternative Identity Splits

Relative to Module D, TriCasRank produced positive SCC differences under all four grouping protocols. The paired differences were 0.080 (95% interval, 0.072–0.090) for target grouping, 0.083 (0.069–0.096) for guide grouping, 0.004 (−0.014–0.027) for atomic-template grouping, and 0.016 (0.001–0.034) for joint grouping. Thus, the improvement was clear under target and guide isolation but small or uncertain under the stricter template-based protocols. Among 1,107 eligible guides, median within-guide SCC increased from 0.696 for Module D to 0.771 for TriCasRank.

Absolute-error performance varied across protocols. TriCasRank reduced MAE relative to Module D under target and guide grouping, showed a slightly higher MAE under atomic-template grouping (0.419 versus 0.415), and a slightly lower MAE under joint grouping (0.413 versus 0.419). The sequence-plus-mapping XGBoost model achieved higher SCC and lower MAE than TriCasRank under both atomic-template and joint grouping. Because these protocols contained only 21 and 20 independent groups, respectively, the results should be treated as boundary-level evidence rather than as stable estimates of performance under identity shift.

### 4.5 Comparison With EasyDesign and Other Approaches

On the matched cohort, TriCasRank outperformed the locally retrained EasyDesign regressor in both ranking and absolute-error metrics. TriCasRank achieved an SCC of 0.809 and an RMSE of 0.363, compared with 0.635 and 0.493, respectively, for EasyDesign-adapted. It also produced a lower absolute error for 67.4% of paired records and had lower MAE across the five observed-activity quintiles (Fig. 4).

This comparison controls for the evaluated records and fold assignments, but it should not be interpreted as a direct comparison with the performance reported in the original EasyDesign publication. EasyDesign-adapted was retrained locally on the shared cohort, whereas the published EasyDesign scores were obtained using different evaluation sets and protocols. The matched analysis therefore compares the two implementations under the present study design rather than establishing superiority over every published EasyDesign setting [2].

Published Cas12a activity predictors have used sequence, mismatch, and positional representations, while EasyDesign addressed guide selection for diagnostic assays and DeepCpf1 modeled guide activity for cellular editing [2,3]. Because these studies differ in assay type, prediction target, data distribution, and validation protocol, their reported scores are not directly comparable with the present results. The matched EasyDesign analysis was therefore used as the primary model comparison.

TriCasRank differs from these approaches mainly in its use of reference-library mapping and training-derived guide and template histories. This design is useful when related identities are represented in the training data, but it also makes performance dependent on the availability and quality of that contextual information. A prospective comparison on an independently collected cohort will be needed to determine whether the observed advantage persists when preprocessing, model selection, and calibration are fixed before evaluation.

### 4.6 Limitations and Future Evaluation

Several limitations temper the interpretation of these results. First, because the historical fixed-validation set was inspected during model development, it cannot be considered a fully independent test set. Similarly, the primary fixed-weight TriCasRank configuration was chosen using target-grouped out-of-fold predictions, meaning its reported performance represents a development-stage estimate. To mitigate selection optimism, a separate nested cross-validation analysis restricted model and weight selection to the inner folds; this process selected the sequence-plus-mapping XGBoost model in 37 of 40 outer-fold evaluations and TriCasRank in the remaining 3. While these findings alleviate some concerns and indicate that TriCasRank was not universally preferred, prospective validation on an independent cohort remains necessary.

Second, relational context was common but not universal in the primary 8,417-record benchmark. Both guide and template-key support were available for 7,531 records (89.5%), whereas 60 records (0.7%) had neither; the remaining records had only one supported key (Fig. 2g). The resulting support gradient, together with the Module D fallback, shows that TriCasRank obtained its strongest gains when relational evidence was available while preserving prediction coverage when that evidence was sparse (Fig. 2a,g,h). The alternative identity splits provide stricter boundary tests but do not replace prospective validation on independent guides, templates and assay batches (Fig. 4a–g).

Third, the source data were derived from an in vitro fluorescence endpoint assay. TriCasRank was not evaluated for cellular editing efficiency, complete kinetic trajectories, other Cas12a orthologs, or substantially different laboratory conditions. The label transformation, replicate structure, plate effects, and raw fluorescence trajectories were also not fully available in the processed dataset. These limitations restrict both biological interpretation and assessment of calibration.

Future evaluation should use a prospectively collected LbCas12a cohort with predefined target-, guide-, and template-level splits. Data processing, feature construction, ensemble weights, and calibration should be fixed before the test data are examined. Recording raw fluorescence trajectories, replicate and plate identifiers, and experimental conditions would also help distinguish model error from assay variation and support evaluation of calibration at the activity extremes.

## 5. Future Work

Our future work will first focus on evaluating generalization to identities and experimental conditions not represented during model development. Although guide- and template-grouped analyses provided stricter separation than the primary target-grouped evaluation, they used the same underlying dataset and do not replace an independent test cohort [11]. A prospective evaluation should therefore define guide-, template-, and target-level partitions before model fitting and keep the final test data unavailable during feature selection, weight optimization, and calibration [12].

We also plan to evaluate physically motivated variables, including predicted hybridization free energy, secondary-structure measures, and position-specific mismatch-energy estimates [13–15]. Their contribution should be tested through predefined feature-block ablations under unchanged data partitions. In the present study, a separate nested cross-validation analysis selected the model and ensemble parameters within the inner folds and evaluated the selected configuration on the outer folds, reducing optimism caused by reuse of the same out-of-fold predictions [12]. Future studies should retain this nested procedure and preregister the model-search space.

The next useful test is a prospective wet-lab screen. Candidate crRNAs should be ranked before synthesis, after which the study can measure the top-ranked hit rate, enrichment over Module D, and correlation with fluorescence. These outcomes would show whether the retrospective improvement actually reduces the number of candidates sent to the laboratory. If the work is extended to diagnostic accuracy, the protocol and report should follow the relevant STARD recommendations [16].

The mapping procedure also needs explicit rules for pathogen variants, IUPAC ambiguity, multiple compatible templates, and unseen template families. A later multi-laboratory study should use versioned models and preprocessing code, preserve its group assignments, and record the predictions produced by each model version. These records would make the analysis reproducible and easier to audit [17].

## 6. Conclusion

In this study, we developed a machine-learning system to rank crRNA–target pairs for Cas12a nucleic acid detection. TriCasRank combines three modules: a sequence-only ensemble (D) and two context-aware modules that use reference mapping and training-derived historical information (B and C). In target-grouped out-of-fold evaluation, TriCasRank achieved an SCC of 0.821, compared with 0.741 for Module D. Its performance was comparable to that of the sequence-plus-mapping XGBoost model, which achieved an SCC of 0.822 and was selected in 37 of 40 nested outer-fold evaluations. On the 2,217-record historical fixed-validation set, the corresponding SCC and PCC were 0.8462 and 0.8355. Because this fixed set had been inspected during model development, these values should be interpreted as historical evaluation results rather than performance on an untouched external cohort.

The results suggest that mapping and historical context can improve candidate ranking when related guide and template identities are represented in the training data. Module D remains important both as part of the final fusion and as a fallback when contextual information is unavailable. However, prediction errors were larger at the extremes of the activity range, and performance was less stable under stricter identity separation. TriCasRank should therefore be treated as a candidate-prioritization tool within the evaluated EasyDesign-derived domain. Whether it can reduce experimental screening, reagent use, or development time must be established through prospective fluorescence assays on independently collected candidates.

## Supporting information

Supplementary Table S6 (fold and subgroup audit)

## Declarations

### Ethics approval and consent to participate

Not applicable.

### Consent for publication

Not applicable.

### Availability of data and materials

Source code, model definitions, predefined fold assignments, out-of-fold predictions, evaluation outputs, bootstrap scripts, and serialized model artifacts associated with this study are available at https://github.com/ZebraL415/Cas12a-Activity-Ranker/releases/tag/paper-2026-10-27

The assay records analyzed in this study were derived from the EasyDesign study [2]. The repository includes SHA-256 checksums for data-provenance verification and the artifacts used to reproduce the reported target-grouped and fixed-set analyses. Access to and reuse of the original assay data remain subject to the conditions specified by the source publication.

### Authors’ contributions

TD conceived and led the study, proposed the controlled comparison and broader evaluation experiments, and coordinated the four-member team’s work, including task allocation and review. TD contributed to feature engineering, model development, evaluation, data visualization and the reproducible codebase, wrote the original manuscript, and coordinated its revision. ZL contributed to model development, implemented fixed-recipe and nested grouped evaluation, and conducted model-comparison, ablation, perturbation and prediction-error analyses. ZL also prepared output figures and source tables, organized the reproducibility code, model artifacts and public release, and drafted and revised the output-related Results and Discussion text and methodological descriptions. HL contributed to conceptualization, data curation and validation, formal statistical analysis, methodology and the evaluation framework, and results generation and verification. SX contributed to data curation and validation; formal analysis of unseen mapping categories, rare identities and fold-level statistics; and review and verification of data partitioning and model evaluation procedures.

## Acknowledgements

Not applicable.

