## Supplementary Tables for "A Three-View System for CRISPR-Cas12a crRNA Activity Ranking: When to Ensemble, When Context Matters More"

The tables below provide the registered supporting analyses, data checks, and comparison context cited in the main text.

### Table S1. Model and evidence registry.

| Model | Version | Input | Protocol | N | SCC | RMSE | Use |
|---|---|---|---|---|---|---|---|
| XGBoost baseline | ranker baseline | 183 active sequence/alignment features | baseline_validation; historical fixed validation | 2217 | 0.768 | 0.423 | Historical early reference comparator |
| XGBoost–CatBoost exploratory blend | ranker final-four exploratory | 183 active sequence/alignment features | baseline_validation; historical fixed validation | 2217 | 0.769 | 0.424 | Historical sensitivity or early ensemble comparison |
| Sequence ensemble A | pre-v2 intermediate | Sequence/pair/guide views without Table S2 mapping | baseline_validation; historical fixed validation | 2217 | 0.797 | 0.399 | Development milestone before mapping-aware system |
| Legacy ABC | previous A+B+C | 1191-context matrix for B/C plus earlier A | baseline_train; target-grouped 5-fold cross-fitted OOF | 8417 | 0.819 | 0.378 | Controlled comparator for release replacement |
| TriCasRank | final system | 183-feature D plus 1191-feature B/C context | baseline_train; target-grouped 5-fold cross-fitted OOF | 8417 | 0.821 | 0.378 | Reproducibility/reference cross-check |

| Model | Version | Input | Protocol | N | SCC | RMSE | Use |
|---|---|---|---|---|---|---|---|
| TriCasRank | evaluation reconstruction | 183-feature D plus 1191-feature B/C context | baseline_train; target-grouped 5-fold paper evaluation reconstruction | 8417 | 0.821 | 0.378 | Main target-isolated endpoint |
| Legacy ABC | previous A+B+C | Historical A plus 1191-context B/C | baseline_validation; historical fixed validation | 2217 | 0.842 | 0.354 | Continuity comparison only |
| TriCasRank | final system | 183-feature D plus 1191-feature B/C context | baseline_validation; historical fixed validation | 2217 | 0.846 | 0.352 | Supplementary release continuity result |
| TriCasRank | evaluation reconstruction | Common-context subset of 183-feature D plus 1191-feature B/C context | baseline_train; target-common-context cross-fitted reconstruction | 5995 | 0.809 | 0.363 | Matched common-context comparison only |

### Table S2. Multiplicity-controlled SCC comparison families.

| Family | Protocol | Comparison | ΔSCC (95% CI) | N | Groups | Holm-adjusted P | Permutations | Scope |
|---|---|---|---|---|---|---|---|---|
| Core (16) | Target | TriCasRank − D | 0.080 (0.072, 0.090) | 8417 | 6825 | 0.008 | 2000 | Paired group-level prediction-swap test |

| Family | Protocol | Comparison | ΔSCC (95% CI) | N | Groups | Holm-adjusted P | Permutations | Scope |
|---|---|---|---|---|---|---|---|---|
| Core (16) | Target | TriCasRank − History prior | 0.287 (0.271, 0.304) | 8417 | 6825 | 0.008 | 2000 | Paired group-level prediction-swap test |
| Core (16) | Target | TriCasRank − Sequence + mapping XGBoost | -0.001 (-0.007, 0.005) | 8417 | 6825 | 1.000 | 2000 | Paired group-level prediction-swap test |
| Core (16) | Target | TriCasRank − D + B fixed | 0.001 (-0.000, 0.002) | 8417 | 6825 | 0.364 | 2000 | Paired group-level prediction-swap test |
| Core (16) | Guide | TriCasRank − D | 0.083 (0.069, 0.096) | 8417 | 1341 | 0.008 | 2000 | Paired group-level prediction-swap test |
| Core (16) | Guide | TriCasRank − History prior | 0.242 (0.223, 0.261) | 8417 | 1341 | 0.008 | 2000 | Paired group-level prediction-swap test |
| Core (16) | Guide | TriCasRank − Sequence + mapping XGBoost | -0.043 (-0.052, -0.033) | 8417 | 1341 | 0.008 | 2000 | Paired group-level prediction-swap test |
| Core (16) | Guide | TriCasRank − D + B fixed | 0.002 (-0.000, 0.004) | 8417 | 1341 | 0.411 | 2000 | Paired group-level prediction-swap test |

| Family | Protocol | Comparison | ΔSCC (95% CI) | N | Groups | Holm-adjusted P | Permutations | Scope |
|---|---|---|---|---|---|---|---|---|
| Core (16) | Template | TriCasRank − D | 0.004 (-0.014, 0.027) | 8417 | 21 | 1.000 | 2000 | Paired group-level prediction-swap test |
| Core (16) | Template | TriCasRank − History prior | 0.636 (0.482, 0.785) | 8417 | 21 | 0.008 | 2000 | Paired group-level prediction-swap test |
| Core (16) | Template | TriCasRank − Sequence + mapping XGBoost | -0.050 (-0.068, -0.024) | 8417 | 21 | 0.040 | 2000 | Paired group-level prediction-swap test |
| Core (16) | Template | TriCasRank − D + B fixed | -0.006 (-0.012, 0.000) | 8417 | 21 | 0.411 | 2000 | Paired group-level prediction-swap test |
| Core (16) | Joint | TriCasRank − D | 0.016 (0.001, 0.034) | 8417 | 20 | 0.472 | 2000 | Paired group-level prediction-swap test |
| Core (16) | Joint | TriCasRank − History prior | 0.702 (0.591, 0.809) | 8417 | 20 | 0.008 | 2000 | Paired group-level prediction-swap test |
| Core (16) | Joint | TriCasRank − Sequence + mapping XGBoost | -0.043 (-0.056, -0.026) | 8417 | 20 | 0.009 | 2000 | Paired group-level prediction-swap test |

| Family | Protocol | Comparison | ΔSCC (95% CI) | N | Groups | Holm-adjusted P | Permutations | Scope |
|---|---|---|---|---|---|---|---|---|
| Core (16) | Joint | TriCasRank − D + B fixed | -0.001 (-0.004, 0.002) | 8417 | 20 | 1.000 | 2000 | Paired group-level prediction-swap test |
| EasyDesign (4) | Target | TriCasRank − EasyDesign-adapted | 0.173 (0.159, 0.188) | 5995 | 5923 | 0.002 | 2000 | Paired group-level prediction-swap test |
| EasyDesign (4) | Guide | TriCasRank − EasyDesign-adapted | 0.162 (0.142, 0.184) | 5995 | 1304 | 0.002 | 2000 | Paired group-level prediction-swap test |
| EasyDesign (4) | Template | TriCasRank − EasyDesign-adapted | 0.073 (0.049, 0.090) | 5995 | 20 | 0.005 | 2000 | Paired group-level prediction-swap test |
| EasyDesign (4) | Joint | TriCasRank − EasyDesign-adapted | 0.055 (0.028, 0.082) | 5995 | 19 | 0.047 | 2000 | Paired group-level prediction-swap test |

**Table S3. Perturbation consistency across predefined seeds.**

| Control | Seed | Unique variable | Delta SCC | Delta PCC | Delta RMSE | Delta MAE | 95% intervals | Status |
|---|---|---|---|---|---|---|---|---|
| history-label shuffle | 137 | Outer-train history-label vector permuted within each fold; sequences, fold IDs, label-free mapping features and true holdout labels unchanged | -0.017 | -0.020 | 0.027 | 0.029 | SCC [−0.022555,−0.010796]; PCC [−0.025812,−0.014100]; RMSE [0.021636,0.031811]; MAE [0.024796,0.032480]; R2 [−0.059088,−0.040922] | Complete; all assertions passed |
| history-label shuffle | 271 | Outer-train history-label vector permuted within each fold; sequences, fold IDs, label-free mapping features and true holdout labels unchanged | -0.019 | -0.022 | 0.028 | 0.029 | SCC [−0.024139,−0.012596]; PCC [−0.027836,−0.015601]; RMSE [0.022913,0.033215]; MAE [0.025646,0.033171]; R2 [−0.061945,−0.043234] | Complete; all assertions passed |
| history-label shuffle | 409 | Outer-train history-label vector permuted within each fold; sequences, fold IDs, label-free mapping features and true holdout labels unchanged | -0.017 | -0.021 | 0.027 | 0.028 | SCC [−0.022671,−0.011435]; PCC [−0.027372,−0.015266]; RMSE [0.022018,0.032286]; MAE [0.024607,0.032198]; R2 [−0.059888,−0.041849] | Complete; all assertions passed |

| Control | Seed | Unique variable | Delta SCC | Delta PCC | Delta RMSE | Delta MAE | 95% intervals | Status |
|---|---|---|---|---|---|---|---|---|
| mapping-block permutation | 137 | Whole mapping block permuted as a block within train and holdout rows; sequence, true labels, folds and XGBoost capacity unchanged | -0.068 | -0.084 | 0.064 | 0.046 | SCC [−0.077626,−0.059672]; PCC [−0.094754,−0.073411]; RMSE [0.056921,0.071230]; MAE [0.041001,0.050629]; R2 [−0.140553,−0.111959] | Complete; all assertions passed |
| mapping-block permutation | 271 | Whole mapping block permuted as a block within train and holdout rows; sequence, true labels, folds and XGBoost capacity unchanged | -0.068 | -0.084 | 0.064 | 0.046 | SCC [−0.077899,−0.060067]; PCC [−0.095521,−0.073882]; RMSE [0.056982,0.071515]; MAE [0.041065,0.050812]; R2 [−0.141231,−0.111898] | Complete; all assertions passed |
| mapping-block permutation | 409 | Whole mapping block permuted as a block within train and holdout rows; sequence, true labels, folds and XGBoost capacity unchanged | -0.068 | -0.084 | 0.064 | 0.045 | SCC [−0.077435,−0.059963]; PCC [−0.095298,−0.073892]; RMSE [0.057016,0.071403]; MAE [0.040849,0.050453]; R2 [−0.141198,−0.112252] | Complete; all assertions passed |

**Table S4. EasyDesign measurement and protocol audit.**

| Topic | Status | Verified finding | Manuscript wording | Excluded inference | Source anchor |
|---|---|---|---|---|---|
| Assay endpoint | SUPPORTED | Fluorescence from an LbCas12a-mediated diagnostic cleavage assay; not genome-editing efficiency. | EasyDesign supplies Cas12a diagnostic fluorescence activity labels. | Do not call the label editing efficiency or clinical diagnostic sensitivity. | ED-M01; Methods: Quantification of Cas12a cleavage activity |
| Wet-lab conditions relevant to label provenance | SUPPORTED | Reported reaction includes 200 ng purified LbCas12a, 25 pM ssDNA FQ probe, 1 μM crRNA, 2 μL sample, 20 μL total at 37 °C. | Use as source-assay provenance only. | Do not claim independent reproduction of every wet-lab condition. | ED-M02; Methods: Quantification of Cas12a cleavage activity |
| 30-min readout | SUPPORTED | Fluorescence was read every 2 min for 2 h; the 30-min value was selected as the rapid activity indicator. | The source activity readout is a 30-min fluorescence-derived value. | Do not infer a kinetic rate from a single 30-min column. | ED-M03; Results/Methods measurement-time passages |
| Plate-control normalization | SUPPORTED | Author-described normalization subtracts the mean of two negative-control wells and divides by the mean of two positive-control wells on the same 96-well plate. | Report the author-described plate-control normalization formula. | Do not say the project recomputed it from raw wells without plate-level data. | ED-M04; Methods: Predicting detection activity: Data processing |
| Time-folding augmentation | SUPPORTED | Published augmentation formula is g(t1,t2)=[[f(t1+2)-f(t1-2)]^((t1-t2)/4)]×f(t2). | The source paper reports time-folding augmentation. | Do not reinterpret this formula as a log or rate transform. | ED-M05; Methods: Data augmentation |

| Topic | Status | Verified finding | Manuscript wording | Excluded inference | Source anchor |
|---|---|---|---|---|---|
| Negative activity-label columns | SUPPORTED | Local Training 30-min and 20-min normalized values are negative; Augment out_logk_measurement is also negative. | The supplied training/augmentation activity-related columns are negative. | Do not infer log base, shift, scale, clamp or physical meaning from negativity. | ED-M06/ED-M07; local workbook value audit |
| Exact numerical label transform | NOT_REPORTED | Exact log base, pseudocount, shift, scale, clamp and inverse transform are not reported. | Use NOT_REPORTED for label-generation details. | Do not write ln, log10, shifted, clipped or inverse-transformed without evidence. | ED-M09; full Methods, local SI and official repository scan |
| Shift/clamp gate | NOT_FOUND_AFTER_SEARCH | No explicit shift formula or clamp/truncation rule was found in the searched sources. | Shift/clamp is unavailable in the supplied sources. | Do not claim a hidden preprocessing gate. | ED-M16; full XML, local SI and official repository scan |
| Dataset count | SUPPORTED | Paper reports 11,496 guide–target pairs; local workbook has 10,634 Training rows and 1,358 Test rows. | State the local workbook count with its source; preserve the paper count separately. | Do not report the paper's 862-test statement as the sole split without qualification. | ED-M10; Methods: Data set preparation; local Table S2 workbook |
| Train/test count conflict | SUPPORTED | A Results paragraph reports 10,634 train + 862 test, whereas Methods and local Test sheet report 1,358 test; the source is internally inconsistent. | Flag the 862 versus 1,358 conflict explicitly. | Do not silently harmonize the numbers. | ED-M11; Results: Development; Methods; local Table S2 |

| Topic | Status | Verified finding | Manuscript wording | Excluded inference | Source anchor |
|---|---|---|---|---|---|
| Split protocol | SUPPORTED | Methods describe completely non-overlapping sets and 20 distinct DNA classes in training for an unseen-target/zero-shot framing. | Attribute non-overlap and 20 training DNA classes to the source paper. | Do not infer an exact random seed or reconstruct the split from the statement alone. | ED-M12; Methods: Data set preparation |
| Augmentation size and gate | SUPPORTED | Augmented dataset reported as 31,993 guide–target pairs; >4-mismatch synthetic pairs were labelled inactive in the CNN12ae description. | Report source-reported augmentation and mismatch gate. | Do not invent mismatch-position distributions or a numeric clamp. | ED-M13/ED-M14; Results: Development; CNN model Methods |
| Classification gate | SUPPORTED | Source evaluates active/inactive using true activity = 3 and reports CNN12ae AUC = 0.8247; this is source-specific. | Attribute the activity=3 gate to the source paper only. | Do not transfer threshold 3 to the project's negative labels without a scale bridge. | ED-M15; Fig. 2 caption and Results: Development |
| Input encoding | SUPPORTED | Model input is 45-position paired target–guide context with 10-dimensional one-hot vectors; '-' denotes insertion/deletion. | Use 45×10 and the source-defined TTTN/21-nt guide context as the input contract. | Do not claim raw flanks where the adapted table omits them. | ED-M17; Methods: One-hot (2D) encoding |

| Topic | Status | Verified finding | Manuscript wording | Excluded inference | Source anchor |
|---|---|---|---|---|---|
| Plate/batch/well mapping | NOT_REPORTED | Plate IDs, batch/run IDs, well coordinates and record-to-plate mapping were not reported in supplied sources. | Treat plate/batch/well covariates as unavailable. | Do not claim plate-level leakage was ruled out. | ED-M19; full XML, local SI and repository scan |
| Control wells and replicates | SUPPORTED + NOT_REPORTED | Two negative-control wells and two duplicate high-fluorescence positive-control wells are reported per plate; sample biological/technical replicate counts are not reported. | Call these control wells; call project rows assay records. | Do not call control duplicates or project rows biological replicates. | ED-M18/ED-M20; Methods and repository audit |
| Raw fluorescence time series | NOT_FOUND_AFTER_SEARCH | Searched package exposes processed spreadsheets but not raw fluorescence time series with plate/well metadata. | Raw plate-level measurements were not found in the searched package. | Do not equate processed tables with public raw measurements. | ED-M27; Data Availability, local SI and official repository scan |
| LbCas12a / TTTN scope | SUPPORTED | Source discussion limits evidence to LbCas12a and PAM TTTN; variants and other Cas12 proteins require further investigation. | Keep project generalization within LbCas12a/TTTN scope. | Do not claim universal guide activity across Cas12 variants. | ED-M25; Discussion |

| Topic | Status | Verified finding | Manuscript wording | Excluded inference | Source anchor |
|---|---|---|---|---|---|
| Project inheritance boundary | SUPPORTED | Project may inherit supplied processed labels and source-defined sequence/task scope, but not undocumented plate mapping or label transform as independently reimplemented. | Use an inherited-versus-unverified data contract. | Do not claim original wet-lab preprocessing was independently reproduced. | ED-M28; evidence synthesis from ED-M01–ED-M27 |

8 **Table S5. Public method-comparison context.**

| Method | Task | Endpoint | Input context | Dataset and validation | Reported evidence | Comparison role |
|---|---|---|---|---|---|---|
| TriCasRank (project snapshot) | EasyDesign Cas12a setting; continuous diagnostic activity | Fluorescence-derived continuous activity | 183-feature D plus 1,191-feature B/C relational context; fixed 0.20D + 0.47B + 0.33C | 8,417 records; 6,825 target groups; Target-grouped five-fold paper evaluation reconstruction | Project endpoint; SCC/PCC/RMSE/MAE/R2 reported in Tables 3 and 5 and Fig. 4 | Project reference; not a published external comparator |
| EasyDesign (CNN12ae) (2024) | LbCas12a; diagnostic crRNA activity ranking | 30-min fluorescence-derived activity with source augmentation | 45-position paired target–guide one-hot encoding; CNN | 11,496 source pairs; 31,993 augmented pairs reported; Source-described non-overlapping target sets; cited test size is inconsistent between 862 and 1,358 | Author reports Spearman rho=0.673 on the cited test and 0.812 on a larger high-mismatch evaluation | Author-reported source protocol; paired local EasyDesign-adapted result is reported separately in Fig. 4i,j |
| ActivityNN (2025) | LbCas12a; diagnostic trans-cleavage activity | Initial 10-min fluorescence slope (RFU/min) | 28 base features + GC + 196 MD/predicted DNA-interaction features | 1,448 targets; separate experimental n=15; Random 1,159/289 split; separate experimental validation n=15 | Author reports ActivityNN test R2=0.9455 (n=289); One-step ActivityNN Pearson r=0.9328 in a separate 15-sample experimental validation; XGBoost R2=0.7777 | Different endpoint, label scale and random split; no same-axis score |

| Method | Task | Endpoint | Input context | Dataset and validation | Reported evidence | Comparison role |
|---|---|---|---|---|---|---|
| Cas12a_predictor (2023) | Cas12a; cellular guide-editing efficiency | Cellular editing efficiency / Pearson validation | Random-forest sequence features, nucleotide bias and mismatches; 4 bp downstream + 26 bp upstream of PAM | Mini-human and endogenous target-site cohorts; external cell datasets; Five-fold CV plus HEK293T-plasmid, HCT116-plasmid and HEK293T-lentivirus validation | Authors report 14–50% relative improvement over compared algorithms; exact coefficients not textually recoverable here | Different cellular endpoint, cohorts and label scale; no project ranking |
| DeepCas12a (2026) | AsCas12a; cellular on-target editing | Binary high/low cellular editing activity from indel frequency | 34-bp target context + DNA methylation + DNase-seq accessibility; CNN + ViT | HT1 N=16,292; external HT2 n=2,963 and HT3 n=1,251; HT1 held-out plus independent external HT2/HT3 | HT1 rho=0.630, r=0.631, AUC=0.868, AP=0.783; external values retained in literature registry | Different enzyme, cellular label and multimodal context; no project ranking |
| ADAPT (2022) | Cas13a; activity-informed viral diagnostic design | Guide activity/classification and design coverage | Pretrained guide-target model plus diversity, specificity and primer-design layers | About 19,000 guide-target training pairs; 1,933 vertebrate-infecting viral species; Author-reported design validation and in-silico coverage; no project-matched target-fold score | Design coverage and time-to-design evidence; not a continuous activity metric | Different enzyme and deployment endpoint; context only, no project ranking |

### Table S6. Electronic fold and subgroup audit workbook.

| Workbook sheet | Rows | Contents |
|---|---|---|
| Fold metrics | 35 | Fold-level metrics for the registered models |
| History support | 4 | Guide and template history support strata |
| Template support | 5 | Candidate-template support strata |
| Activity bins | 80 | Calibration and error summaries across activity bins |
| Guide ranking | 4 | Within-guide ranking summaries |
| Provenance | 6 | Source and derivation registry for the supplementary workbook |
